## Supplementary Figure 1 and Tables 1, 2 for "Spinal neural tube formation and tail development in human embryos"

**Santos et al - Supplementary Information**

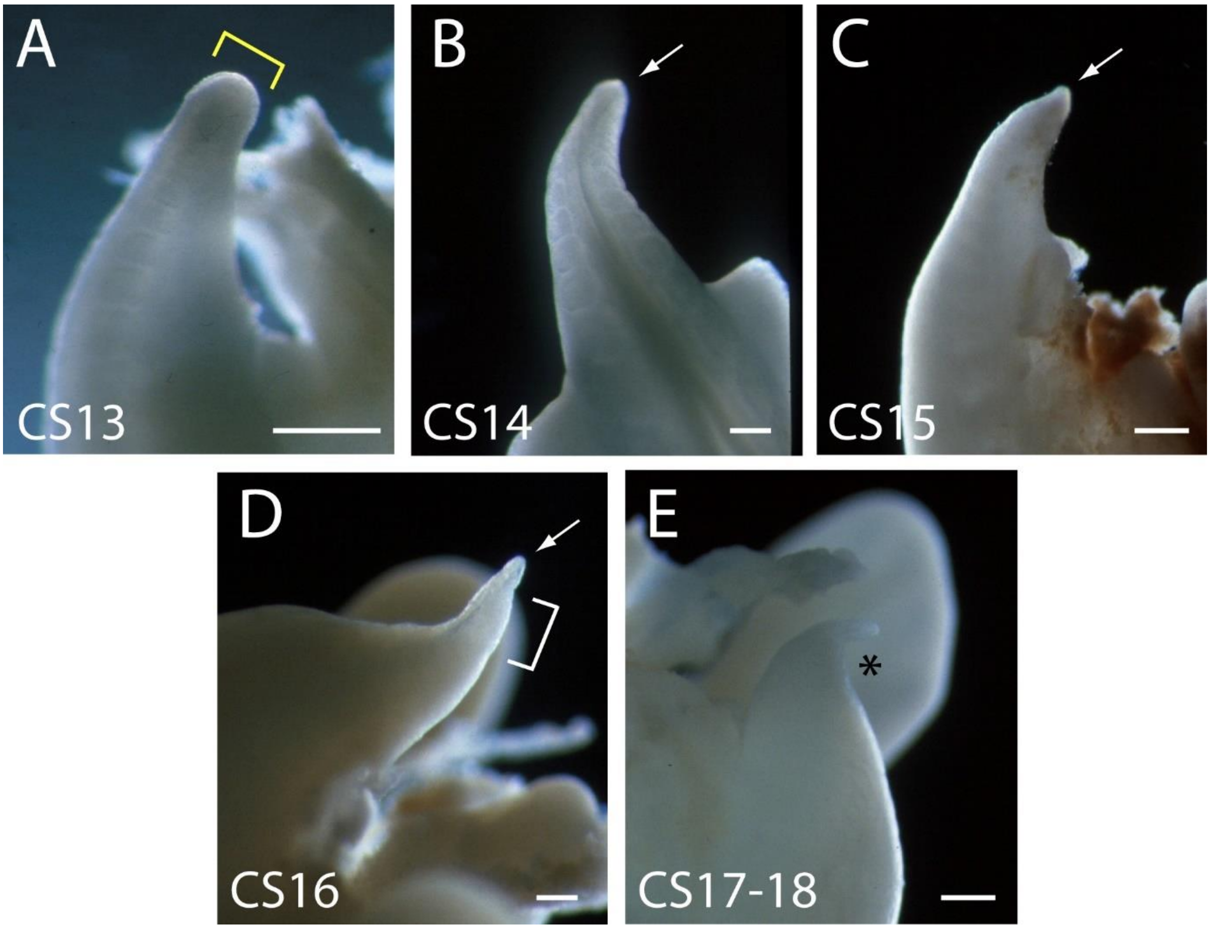


**Supplementary Figure 1.** Human embryonic tails (additional to those in Figure 2) to show the reproducibility of tail morphology changes between CS13 and CS18. Note the broad tailbud at CS13 with rounded tip (yellow bracket in A), which gradually narrows and becomes increasingly pointed, through CS14, CS15 and CS16 (arrows in B,C,D). At CS16, the tail tip has a markedly translucent appearance (bracket in D). By CS17-18, the tail has shortened and is dorsally curved at its tip (asterisk in E). Scale bars: 0.5 mm.

**Supplementary Table 1. Publications (n = 28) on human secondary neural tube and body formation, showing source and number of embryos (n = 925 total) and the topics covered**

| Study | Samples | PNP closure | Primary to secondary NT transition | Mode of secondary NT formation | Multiple NT lumens observed | Formation of secondary somites, notochord, tail-gut | Morphology of tail regression | Mode of cell death during regression | Gene expression |
|---|---|---|---|---|---|---|---|---|---|
| **Kunitomo** (1918). Carnegie Inst. Contr. Embryol 8, 161-198. | 44 embryos, 4-125 mm crown-rump length, sections analysed from Carnegie collection | | ✓ | ✓ | ✓ | | ✓ | | |
| **Streeter** (1919). Am. J. Anat. 25, 1-11. | Unspecified number of embryos, sections analysed from Carnegie collection | | | ✓ | ✓ | | ✓ | | |
| **Bolli** (1966). Acta. Anat. 64, 48-81. | 123 normal embryos, 2-150 mm CRL, with unspecified method of collection | | | ✓ | ✓ | | ✓ | | |
| **Lemire** (1969). Teratology 2, 361-370. | 8 embryos of CRL 6-25 mm, CS14-21, including from spontaneous abortions (3) and ectopic pregnancies (4) | | | ✓ | ✓ | | ✓ | ✓ | |
| **Hughes** et al. (1974). J. Embryol. Exp. Morphol. 32, 355-363. | 6 embryos at 28-40 days, sections analysed, with unspecified method of collection | | | ✓ | ✓ | | | | |
| **Fallon** et al. (1978). Am. J. Anat. 152, 111-129. | 52 embryos at CS14-22 from induced abortion | | | ✓ | ✓ | | ✓ | ✓ | |
| **Müller** et al. (1987). Anat. Embryol. 176, 413-430. | 24 embryos at CS12, sections analysed from Carnegie collection | ✓ | ✓ | ✓ | | ✓ | | | |
| **Müller** et al. (1988). Anat. Embryol. (Berl) 177, 203-224. | 25 embryos at CS13, sections analysed from Carnegie collection | | | ✓ | ✓ | ✓ | | | |
| **Nievelstein** et al. (1993). Teratology 48, 21-31. | 36 embryos CS11-23 of unspecified origin | ✓ | ✓ | ✓ | | | ✓ | ✓ | |
| **Saraga-Babic** et al. (1994). Ann. Anat. 176, 277-286. | 20 embryos at 4-12 weeks, obtained from artificial abortions | | | ✓ | | ✓ | ✓ | | ✓ |
| **Saraga-Babic** et al. (1995). J. Brain Res. 36, 341-347. | 15 embryos aged 4-8 developmental weeks (CS12-23) from artificial abortions | ✓ | ✓ | ✓ | ✓ | | | | |
| **Peeters** et al. (1998). Anat. Embryol. 198, 185-194. | 11 embryos CS10-12, photographs analysed, of Kyto embryos from induced abortion for social reasons | ✓ | | | | | | | |
| **Nakatsu** et al. (2000). Anat. Embryol. (Berl) 201, 455-466. | 68 normal embryos CS10-12 from pregnancy terminations for social reasons; a few from spontaneous or emergency abortions | ✓ | | | | | | | |
| **Sapunar** et al. (2001). Ann. Anat. 183, 217-222. | 8 embryos at CS12-18 obtained from artificial abortions | | | | | | ✓ | ✓ | |
| **O'Rahilly** et al. (2002). Teratology 65, 162-170. | 98 embryos at CS8-13, sections analysed from Carnegie collection | ✓ | | | | | | | |
| **Müller** et al. (2004). Cells Tissues Organs 177, 2-20. | 52 embryos at CS9-23, sections analysed from Carnegie collection | ✓ | ✓ | ✓ | ✓ | ✓ | ✓ | | |

| | | | | | | | | | |
|---|---|---|---|---|---|---|---|---|---|
| **Saitsu** et al. (2004). Anat. Embryol. (Berl) 209, 107-117. | 20 embryos around stage of posterior neuropore closure (CS12-13). Most from induced abortion for social reasons; one from ectopic pregnancy | ✓ | ✓ | ✓ | ✓ | | | | |
| **Vilovic** et al. (2006). Anat. Embryol. 211, 1-9. | 18 embryos at CS12-CS23 (5-12 weeks), from spontaneous or induced abortions | | | | | | ✓ | ✓ | ✓ |
| **Pytel** et al. (2007). Folia Morphol. (Warsz. ) 66, 104-108. | 12 embryos at CS13-17, 32-41 days, with unspecified method of collection | | | ✓ | ✓ | | | | |
| **Saitsu** et al. (2008). Congenit. Anom. (Kyoto) 48, 1-6. | 43 embryos obtained from Kyoto collection from induced abortion for social reasons | ✓ | ✓ | ✓ | ✓ | | | | |
| **Yi** et al. (2010), FASEB J 24, 3341 3350. | 18 embryos at weeks 4-9 from mifepristone-induced abortions | | | | | | | | ✓ |
| **Fang** et al. (2010), Dev Cell 19, 174-184 | 123 embryos at CS9-14 from induced abortions | | | | | | | | ✓ |
| **Krupp** et al. (2012). Birth Defects Res. A Clin. Mol. Teratol. 94, 683-692. | 8 or more CS12 and CS13 embryos, from mifepristone-induced abortions | | | | | | | | ✓ |
| **Olivera-Martinez** et al. (2012). PLoS Biol .10, e1001415. | 3 embryos at CS12 (2) and CS16 (1) from induced abortion for social reasons | | | | ✓ | | | | ✓ |
| **Yang** et al. (2014). Childs Nerv. Syst. 30, 73-82. | 21 embryos at CS12-23, from therapeutic pregnancy termination or surgical procedures to remove uterus/oviduct | | | ✓ | ✓ | ✓ | ✓ | ✓ | ✓ |
| **Jang** et al. (2016). Pediatr. Neurosurg. 51, 9-19. | 20 embryos and fetuses, 6-14 weeks gestation from miscarriages and ectopic pregnancies | | ✓ | | | | | | |
| **Tojima** et al. (2018). J. Anat. 232, 806-811. | 42 embryos at CS13-23 from Kyoto collection from induced abortion for social reasons | | | | | ✓ | ✓ | | |
| **Xu** et al. (2023). Nat Cell Biol 25, 604 | 7 embryos at CS12-14 (4-6 weeks) from medical terminations of pregnancy | | | | | | | | ✓ |

Abbreviations: CS, Carnegie Stage; NT, neural tube; PNP, posterior neuropore

**Supplementary Table 2. Human embryos as summarised in Figure 2I-K and Table 2 ***

| Carnegie Stage | Days pc | Somite no. | Crown-rump length (mm) ** | Tail length (mm) ** | Tail length distal to somites (mm) ** |
|---|---|---|---|---|---|
| 13 | 28 | 37 | 7.00 | 0.80 | |
| | 28 | 38 | 5.33 | 0.78 | 0.29 |
| | 28 | 34 | 4.66 | 0.69 | 0.50 |
| | 29 | 35 | 6.58 | 0.63 | 0.38 |
| | 30 | 38 | 6.88 | 1.75 | 0.52 |
| | 30 | 34 | 8.16 | 1.30 | 0.47 |
| | 30 | 32 | | 1.45 | 0.76 |
| 14 | 31 | 36 | | | |
| | 31 | 32 | | | |
| | 31 | 36 | 8.70 | 1.15 | 0.60 |
| | 31 | 37 | 6.00 | 1.08 | 0.52 |
| | 32 | 34 | 10.38 | 0.75 | |
| 15 | 33 | 37 | 9.00 | 1.20 | 0.56 |
| | 33 | 34 | 8.60 | 2.00 | 0.91 |
| | 34 | 37 | 9.75 | 1.48 | 0.60 |
| | 34 | 38 | 7.80 | | 0.78 |
| | 35 | 37 | 9.38 | 1.25 | 0.60 |
| | 35 | 34 | | 0.50 | 0.54 |
| | 35 | 34 | | 0.65 | |
| 16 | 37 | 39 | | 1.10 | |
| | 37 | 39 | | 1.05 | 0.35 |
| | 37 | 33 | 12.33 | 1.19 | 0.38 |
| | 37 | 39 | | 0.95 | |
| | 37 | 35 | 12.50 | 2.44 | 0.64 |
| | 38 | 38 | 10.13 | 1.08 | |
| | 38 | 37 | 10.66 | 1.20 | 0.46 |
| | 39 | 33 | 12.75 | 1.33 | |
| 17 | 40 | 34 | 10.88 | 1.44 | 0.35 |
| | 40 | 34 | 13.25 | 1.25 | |
| | 42 | 31 | | 1.05 | 0.33 |
| | 43 | 30 | 12.50 | 1.00 | |
| 18 | 44 | | | | |
| | 44 | 33 | 15.88 | 1.25 | 0.00 |
| | 45 | 31 | 18.13 | 0.63 | |
| | 45 | 34 | 14.50 | 1.38 | |
| | 45 | 33 | 13.00 | 1.76 | |
| | 45 | 32 | | 0.70 | |

* Each line in the table corresponds to a different human embryo (n = 37).

** Measurements of crown-rump length, tail length and tail length distal to somites were not available for all embryos. Data in Figure 2I-K and Table 2 are based on the available data.
